# Visualization of Retroplacental Clear Space Disruption in a Mouse Model of Placental Accreta

**DOI:** 10.1101/2022.07.12.499572

**Authors:** Andrew A. Badachhape, Prajwal Bhandari, Laxman Devkota, Mayank Srivastava, Eric A. Tanifum, Verghese George, Karin A. Fox, Chandrasekhar Yallampalli, Ananth V. Annapragada, Ketan B. Ghaghada

## Abstract

**Introduction:** Prior preclinical studies established the utility of liposomal nanoparticle blood-pool contrast agents in visualizing the retroplacental clear space (RPCS), a marker of normal placentation, while sparing fetuses from exposure because the agent does not cross the placental barrier. In this work we characterized RPCS disruption in a mouse model of placenta accreta spectrum (PAS) using these agents.

**Methods:** Contrast-enhanced MRI (CE-MRI) and computed tomography (CE-CT) using liposomal nanoparticles bearing gadolinium (liposomal-Gd) and iodine were performed in pregnant Gab3^-/-^ and wild type (WT) mice at day 16 of gestation. CE-MRI was performed on a 1T scanner using a 2D T1-weighted sequence (100×100×600 µm^3^ voxels) and CE-CT was performed at a higher resolution (70×70×70 µm^3^ voxels). Animals were euthanized post-imaging and feto-placental units (FPUs) were harvested for histological examination. RPCS conspicuity was scored through blinded assessment of images.

**Results:** Pregnant Gab3^-/-^ mice show elevated rates of complicated pregnancy. Contrast-enhanced imaging demonstrated frank infiltration of the RPCS of Gab3^-/-^ FPUs. RPCS in Gab3^-/-^ FPUs was smaller in volume, demonstrated a heterogeneous signal profile, and received lower conspicuity scores than WT FPUs. Histology confirmed *in vivo* findings and demonstrated staining consistent with a thinner RPCS in Gab3^-/-^ FPUs.

**Discussion:** Imaging of the Gab3^-/-^ mouse model at late gestation with liposomal contrast agents enabled *in vivo* characterization of morphological differences in the RPCS that could cause the observed pregnancy complications. An MRI-based method for visualizing the RPCS would be valuable for early detection of invasive placentation.

## Introduction

Placenta Accreta Spectrum (PAS) describes a range of conditions where pathological invasion of the myometrium by placental tissue complicates delivery and increases the risk for adverse maternal outcomes [1,2]. A compelling theory for this disorder is that surgical procedures such as caesarian delivery, myomectomy, or hysteroscopic resection can cause scarring and disrupt endometrial and myometrial integrity [3]. When placental tissue implants over the scar, the risk for non-separation and hemorrhage during delivery increases due to disruption of the placental-myometrial interface and potential involvement of contiguous visceral structures that may be enveloped in the attendant fibrosis and neovascularization. The incidence of PAS has increased over the past several decades and is responsible for 1.1% of all maternal deaths in the U.S. [4,5].

The most common management of PAS is via cesarean delivery followed by hysterectomy. This approach requires a high level of surgical skill and results in permanent sterility. Alternatively, uterus-sparing approaches may be attempted, either by leaving the placenta in situ after delivery and awaiting resorption or expulsion, or by resecting the area of myometrium affected by pathologic implantation and repairing the uterus as a single-step resection and repair technique. Disadvantages of uterus-sparing approaches include long resolution time (up to 9-months) and the need for delayed hysterectomy should bleeding or infection develop in the case of conservative management; additionally, the potential for development of choriocarcinoma from the residual placenta has also been reported [6]. Focal resection and repair may not be possible in cases of deep cervical and lateral placental extension and requires a margin of healthy tissue superior to the cervix to which the upper margin of resection can be repaired, which can be difficult to discern prior to delivery.

Whenever the abnormal placenta implants low within the uterus, such as in the lower segment or cervix behind the bladder or extends laterally (involving the parametria), the risk for hemorrhage or urinary tract injury is further increased. Major pelvic structures including uterine blood vessels and the ureters course through the parametria near the lower uterus. Significant blood loss requiring transfusion and ICU admission are reduced with accurate antenatal diagnosis of PAS and delivery by experienced multidisciplinary teams. Timely referral to centers that are staffed by experienced clinical teams hence depends directly on the ability to detect PAS antenatally.

Visualization of the uteroplacental interface, or retroplacental clear space (RPCS), is an indicator of normal placentation, but RPCS is difficult to assess with ultrasound and conventional magnetic resonance imaging (MRI) [7–10]—the preferred cross-sectional imaging modalities employed in pregnancy since they do not involve ionizing radiation. Because of the difficulties in diagnosis, the utility of MRI has been debated, with some experts advocating its utility in identifying topography and extent of invasion to optimize surgical planning [11], while others have questioned whether it falsely changes management. While both ultrasound and MRI have a reported sensitivity and specificity approaching or above 90% in expert centers [12–14], the overall rate of antenatal detection in multiple population-based studies is closer to 30-50% [7,15,16]. In cases of shallow or focal invasion, antenatal diagnosis is especially challenging, even in expert centers, highlighting the pressing need for improved antenatal diagnostic imaging.

Further complicating study of the RPCS is the lack of preclinical models of invasive placentation. Recent studies demonstrated the creation of a mouse model of accreta through surgical techniques that damage the uterus either in a non-pregnant mouse or immediately post-partum [17]. Uterine scarring techniques in rodents may provide a strong model for future study of PAS, however these are difficult procedures to perform reproducibly. A mouse model of spontaneous accreta would be exceptionally valuable in this regard, and a promising candidate has emerged in the Gab3 knockout mouse model (Gab3^-/-^), which has been found to have higher rates of dystocia and placental invasion relative to wild type counterparts [18].

We have previously demonstrated a high T1 relaxivity liposomal nanoparticle-based gadolinium (liposomal-Gd) blood-pool contrast agent for contrast-enhanced magnetic resonance imaging (CE-MRI) -based visualization of the RPCS in mouse and rat models [19–21]. In this study, we demonstrate the use of liposomal-Gd CE-MRI to identify disruption of the RPCS in a Gab3-/-mouse model of adherent placentation. CE-MRI findings were confirmed through high-resolution contrast-enhanced computed tomography (CE-CT) and histopathology.

## Materials and Methods

### Animal Model and Breeding Study

All studies involving mice were performed using an animal use protocol approved by the Institutional Animal Care and Use Committee at Baylor College of Medicine. Gab3^*-/-*^ mice were generated through CRIPSR-Cas9 genome editing and breeding pairs were provided by the Cincinnati Children’s Hospital Medical Center (CCHMC) [18]. Gab3^-/-^ female mice were generated through breeding Gab3^-/-^ male mice with Gab3^*+/-*^ female mice. Three pregnant female Gab3^-/-^ mice and three pregnant wild type (WT) mice (8-12 weeks old; ∼20-30 g body weight before pregnancy) were used in the imaging study. All Gab3^-/-^ female mice were impregnated by Gab3^-/-^ male mice and all WT female mice were impregnated by WT male mice. The first day of gestation, designated as 0.5, was when a vaginal copulation plug was detected. Imaging was performed on day 16.5 of pregnancy (E16.5). For the imaging study, 28 Gab3^-/-^ FPUs and 31 WT FPUs were examined.

An additional 3 pregnant female Gab3^-/-^ mice and 3 pregnant WT mice with similar age and weight ranges were used to observe differences in FPU viability during pregnancy in a controlled breeding study. As with the imaging study, all Gab3^-/-^ female mice were impregnated by Gab3^-/-^ male mice and all WT female mice were impregnated by WT male mice. For the breeding viability study, the outcomes of 54 Gab3^-/-^ FPUs and 69 WT FPUs were assessed across 2-3 cycles of breeding for each pair over a 4-month period. A non-contrast MRI was performed at E12.5 to determine the number of FPUs generated during pregnancy, and the dams/pups were monitored for 1 week post-delivery to determine the outcome of FPUs (Table 1).

**Table 1:** Observed pregnancy outcomes and delivery dates for 3 Gab3^-/-^ experimental breeding pairs (Gab EP1, Gab EP2, Gab EP3) and 3 WT experimental breeding pairs (WT EP1, WT EP2, WT EP3) over 3 pregnancy periods (P1, P2, P3). Viable fetoplacental units (FPU) are listed with the total number of FPU in parentheses or listed as “Did Not Deliver” (DND).

|  | P1 |  | P2 |  | P3 |  |
| --- | --- | --- | --- | --- | --- | --- |
|  | Delivery | Viable FPU | Delivery | Viable FPU | Delivery | Viable FPU |
| GAB EP1 | E17.5 | 0 (10) | E18.5 | 0 (6) | DND | DND |
| GAB EP2 | E18.5 | 7 (12) | E18.5 | 0 (7) | DND | DND |
| GAB EP3 | E20.5 | 4 (7) | E17.5 | 0 (6) | E20.5 | 4 (6) |
| WT EP1 | E20.5 | 8 (8) | E20.5 | 6 (6) | E20.5 | 7 (7) |
| WT EP2 | E21.5 | 10 (10) | E19.5 | 5 (7) | E20.5 | 8 (8) |
| WT EP3 | E20.5 | 9 (9) | E20.5 | 5 (8) | E20.5 | 6 (6) |

### Magnetic Resonance Imaging (MRI)

Imaging was performed on a 1T permanent MRI scanner (M2 system, Aspect Imaging, Shoham, Israel) with a 35 mm transmit-receive RF volume coil, similar to methods described previously[19,20,22]. Briefly, animals were sedated using 3% isoflurane, placed on the MRI animal bed, and then maintained at 1-2% isoflurane delivered using a nose cone setup. Animal breathing rate was monitored through pressure pad placed below the animals abdomen.

Liposomal-Gd formulations were prepared as per methods described previously [19,20,22]. This agent exhibits a long half-life (17.5 ±1.5 hours) and T1 relaxivity is 31 ± 5 (*s* ·*mM*)^−1^ [23,24]. Liposomal-Gd was administered intravenously via the tail vein (0.15 mmol Gd/kg).

All animals underwent pre-contrast and post-contrast scans at day 16 of gestion (E16.5). T1-weighted (T1w) scans were acquired using a 2D gradient echo (GRE) sequence. Post-contrast scans were performed ∼5-20 minutes post injection and 1-1.5 hours after pre-scans. Scan parameters for T1w-GRE sequence were echo time (TE) = 8 ms, repetition time (TR) = 32 ms, flip angle = 70 °, slice thickness = 0.6 mm, field of view = 50 mm, number of slices = 30-35, matrix = 500 × 500, acquisition plane = coronal; in-plane resolution = 100 × 100 µm^2^, number of excitations (NEX) = 1, scan time ∼ 10 minutes. T2-weighted scans were acquired with a fast spin echo sequence (T2w-FSE). T2w-FSE scan parameters were: echo time (TE) = 80 ms, repetition time (TR) = 6816 ms, slice thickness = 0.8 mm, field of view = 80 mm, number of slices = 33, matrix = 256 × 250, acquisition plane = coronal; in-plane resolution = 312.5 × 320 µm^2^, NEX = 2, echo train length = 2, scan time ∼ 6 min. The same T1w-GRE and T2w-FSE sequences were used to acquire post-contrast scans. Coil calibrations were repeated between animals at and frequency calibration, transmitter gain adjustment, receiver gain adjustment, and dummy scans were automatically performed before each acquisition.

Three acquisitions were performed for each GRE scan protocol and two acquisitions were performed for each FSE scan protocol. Magnitude-sum averages were generated in Matlab® (version R2015a, MathWorks©, Natick, MA) using acquisitions that showed little to no motion artifact.

### Computed Tomography (CT) imaging

High-resolution, contrast-enhanced CT imaging [19,25] on a small animal micro-CT system (Siemens Inveon) was used for secondary confirmation of MRI findings and to acquire high-resolution isotropic views of the RPCS. Animals were sedated with 3% isoflurane, setup on the CT animal bed, and then maintained at 1-2% isoflurane delivered using a nose cone setup. As with MRI studies, a pressure pad below the animal was used to monitor respiration rate during imaging.

The scan parameters for the CT image acquisition were: 50 kVp, 0.5 mA, 850 ms X-ray exposure, 540 projections, 70 µm isotropic spatial resolution, scan time ∼ 10 minutes. Contrast-enhanced CT imaging was performed after intravenous administration of a liposomal-iodine blood pool contrast agent (1.65 mg I/g body weight) [25–27].

### Histology

Animals were euthanized after imaging and FPUs from each animal were harvested and stored intact in 4% formalin. The FPUs were paraffin-embedded, cut into 8 μm thick sections and stained with Hematoxylin and Eosin (H&E).

### Image Analysis

All MR and CT images were analyzed with OsiriX (version 5.8.5, 64-bit) and MATLAB (version 2015a). Regions-of-interest were manually drawn to segment the placenta (P), amniotic fluid (AF) compartment, and the retroplacental clear space (RPCS) from 2D GRE images. Volumetric and signal data from these ROIs were determined for the segmented structures, and the data is presented in terms of average and standard deviation.

#### Review of MR and Histopathology Images

Two blinded observers with experience in placental imaging reviewed all post-contrast T1w-GRE scans. For each animal at each time point, averaged T1w-GRE images were reviewed for the visualization of RPCS for all FPUs. The visibility of RPCS was scored for each FPU by the observers on a 3-point scale: 0 = not visible, 1 = partly visible, and 2 = clearly visible. A single blinded observer with experience in histological review of placental tissues then rated H&E stained sections obtained from 10 WT FPUs and 9 Gab3^-/-^ FPUs on the same scale. Data is presented as average and standard deviation of the scores.

#### Quantitative Analysis

In additional to volumetric data, contrast-to-noise ratio (CNR) was determined from three post-contrast T1w-GRE acquisitions using methods demonstrated in previous research studies [20]. This calculation considers signal in the placenta (*S*_*p*_) and signal in the retroplacental clear space (*S*_*R*_) relative to background noise (*N*) according to Equation 1:

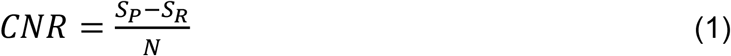

Note that this analysis is the same as finding the difference in signal-to-noise ratio (SNR) for both regions of interest when using a common noise reference.

#### Statistical Analysis

The Wilcoxon rank sum test was used for statistical analysis of placental and retroplacental clear space volumes segmented from the post-contrast GRE images. Fisher’s Exact test was used to confirm statistical significance between Gab3^*-/--*^ and WT FPU outcomes in the breeding study.

## Results

Pregnant Gab3^-/-^ mice demonstrated higher rates of premature delivery and pregnancy complications than wild type (WT) counterparts (Table 1). In addition to giving birth earlier, the fetuses of Gab3^-/-^ pregnant mice were nonviable (stillborn) at delivery at a significantly higher rate than seen in WT pregnant mice (Figure 1e). After two generations of breeding, two Gab3^-/-^ breeding pairs (Gab EP1 and Gab EP 2) were unable to conceive a third generation while all 3 WT breeding pairs successfully delivered a third litter.

**Figure 1:**
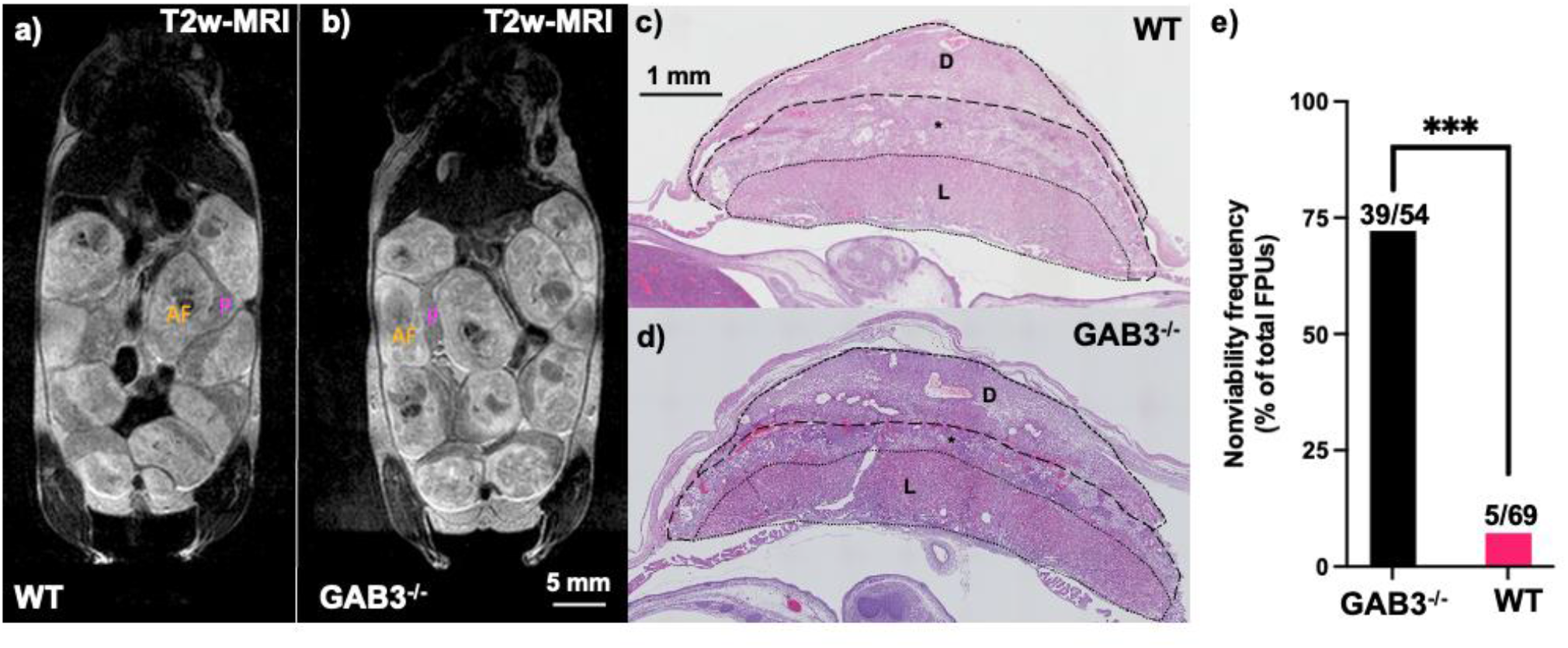
Gab3^-/-^ mice demonstrate complications during pregnancy and disruption across the retroplacental clear space (RPCS) or junctional zone that are not observable *in vivo* with non-contrast MRI. (a-b) Anatomical T2-weighted fast-spin echo magnetic resonance imaging showed no differences in gestational development between Gab3^-/-^ mice and wild type (WT) counterparts at 16 days of gestation. (c-d) Hemaxytolin and Eosin (H&E) staining demonstrates a less heavily stained RPCS, or junctional zone (*), between the placental labyrinth (L) and decidua (D) in WT fetoplacental units relative to Gab3^-/-^ fetoplacental units. e) Gab3^-/-^ pregnant mice demonstrated higher rates of dystocia and premature delivery which led to a greater number of nonviable births relative to WT pregnant mice (Fisher’s exact test, p<0.0005).

MRI studies to investigate potential reproductive anomalies in the Gab3^-/-^ model began with non-contrast T2-weighted anatomical imaging to assess anatomical or structural differences in the placenta or RPCS. Gab3^-/-^ and WT FPUs showed no obvious differences at E16.5 on T2-weighted FSE images, however H&E staining revealed thinner and denser regions within the RPCS (Figure 1a-d) in the Gab3^-/-^ FPUs. In clinical cases, T2-weighted scans of patients diagnosed with PAS often show dark intraplacental bands, abnormal vascularity, or focal thinning of the myometrium [28,29]. None of those features were seen in T2-weighted images acquired in this study. High resolution nanoparticle contrast-enhanced MRI (CE-MRI) demonstrated a heterogeneous and interrupted RPCS in Gab3^-/-^ FPUs relative to WT counterparts (Figure 2). These qualitative differences were confirmed at higher resolution (70 µm isotropic voxels) in nanoparticle CE-CT images (Figure 3).

**Figure 2:**
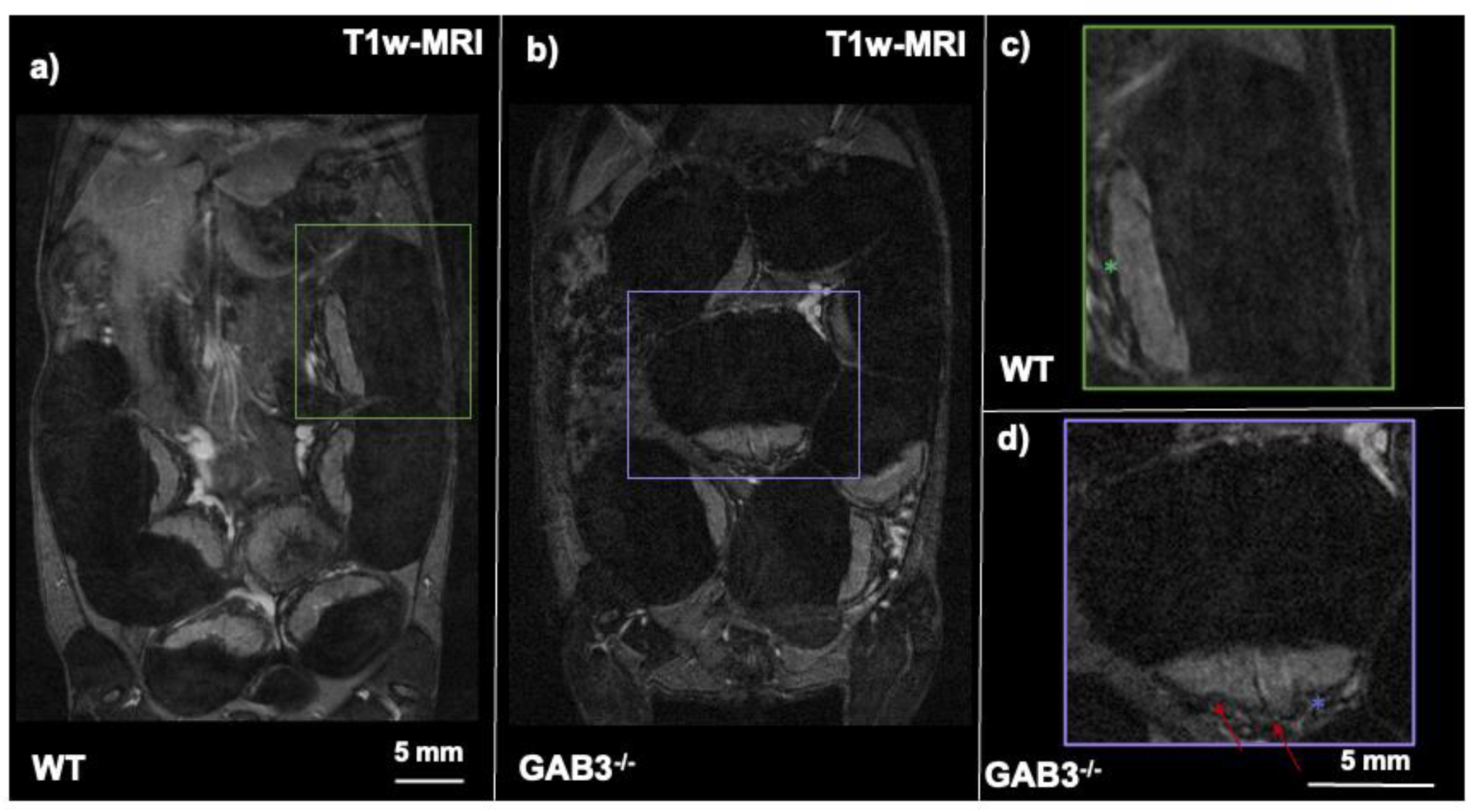
High-resolution contrast-enhanced magnetic resonance imaging (CE-MRI) demonstrates differences in placental margins and the retroplacental clear space (RPCS) in pregnant Gab3^-/-^ and wild type (WT) mice. (a-b) Coronal T1-weighted gradient recalled echo acquisitions for WT and Gab3-/-mice demonstrate enhancement in the placenta post contrast administration with the RPCS visualized as a hypointense region between the placenta and decidua. The RPCS (*) and placental margins appear smoother and darker in WT FPUs (c) and have a more homogeneous and less jagged appearance than in Gab3^-/-^ FPUs (d). Red arrows indicate jagged or obscured regions.

**Figure 3:**
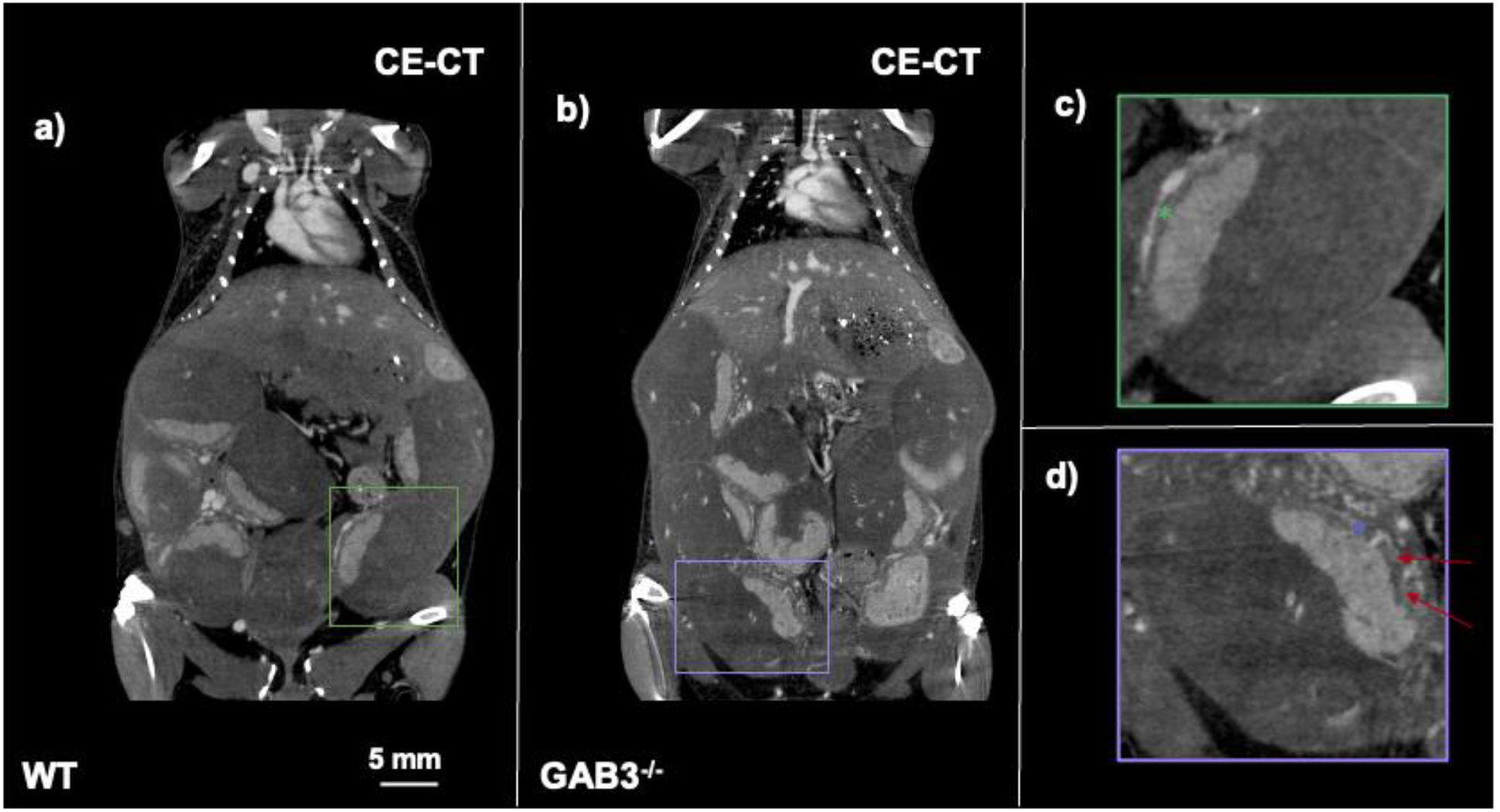
Contrast-enhanced computed tomography (CE-CT) images acquired with 70 um isotropic voxels were used to further confirm in vivo magnetic resonance imaging findings. (a-b) Coronal images of the abdominal region of pregnant wild type and Gab3^-/-^ mice demonstrate enhancement in the placenta relative to a hypointense region denoted as the RPCS. As with T1-weighted MRI, the RPCS (*) in WT FPUs (c) appears wider and easier to identify than in Gab3^-/-^ FPUs (d). Red arrows indicate jagged or obscured regions.

Quantitative assessment of contrast-enhanced images further emphasized differences in the appearance of the RPCS between Gab3^-/-^ and WT FPUs (Figure 4). Blinded review of CE-MR images demonstrated a significant difference (p <0.05) in the mean visibility score of the RPCS in FPUs of Gab3^-/-^ (28 FPUs,1.10 ± 0.48) and Gab3^*+/+*^ (31 FPUs,1.55 ± 0.43) mice. Observations from blinded review identified Gab3^-/-^ RPCS as thinner and more difficult to identify. Gab3^-/-^ RPCS volumes were consistently smaller and appeared with more heterogeneous signature throughout the segmented volume. MRI-derived mean RPCS volume were found to be significantly lower (p<0.005) in Gab3^*/-*^ FPUs (9.7 ± 0.9 mm^3^) than in WT FPUs (12.1 ± 1.8 mm^3^). MRI-derived mean RPCS signal was significantly higher (p<0.005) in Gab3^-/-^ RPCS than in WT RPCS, leading to lower placental-RPCS CNR in Gab3^-/-^ FPUs compared to WT FPUs.

**Figure 4:**
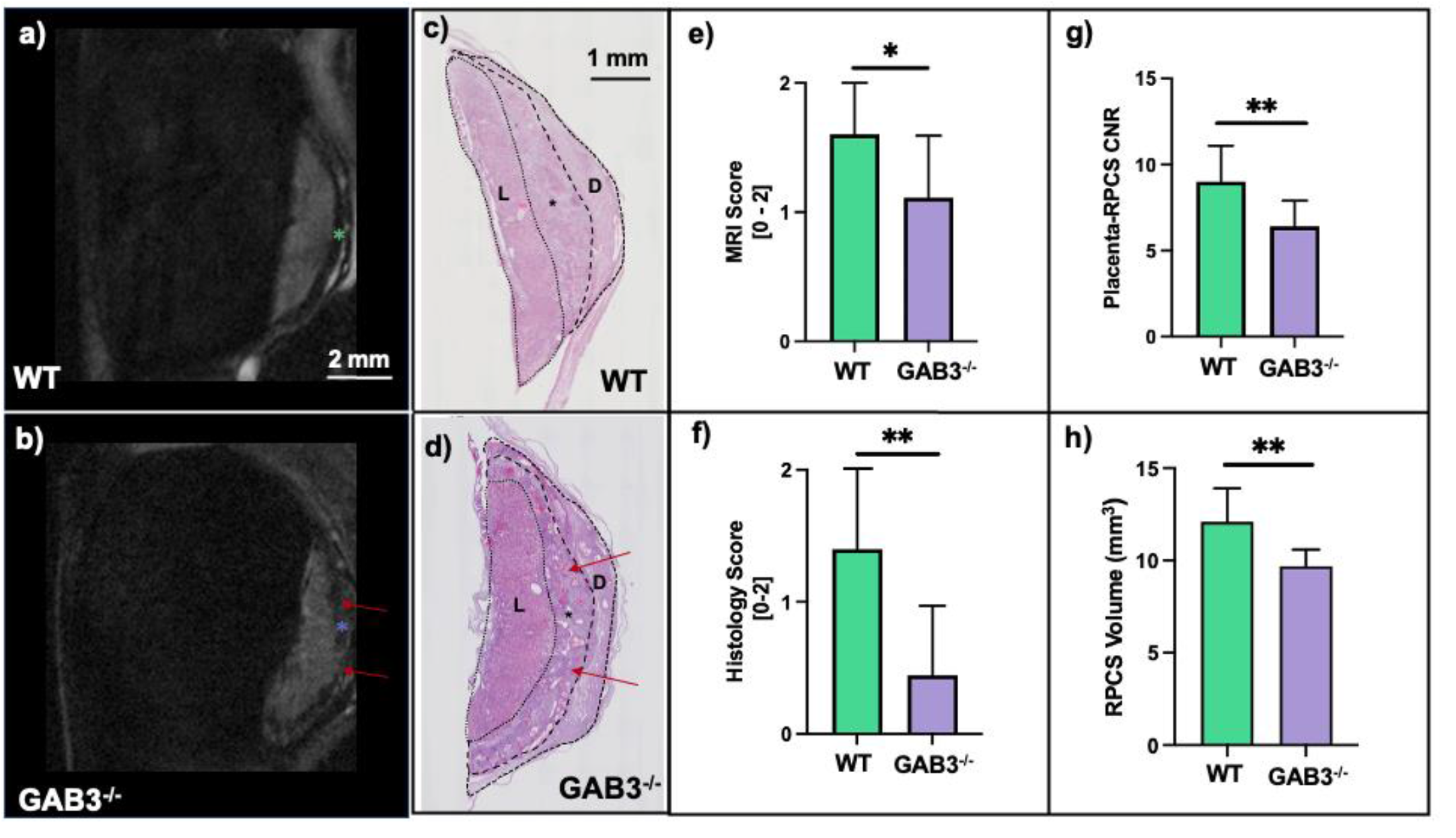
Quantitative assessment of the retroplacental clear space (RPCS) volume further differentiates the appearance between Gab3^-/-^ fetoplacental units (FPUs) and wild type (WT) FPUs. Comparison of T1-weighted magnetic resonance (MR) images of a) WT and b) Gab3^-/-^ FPUs with ex vivo hematoyxlin and eosin (H&E) of c) WT and d) Gab3^-/-^ FPUs demonstrate a thinner and more heterogeneous RPCS (*) in Gab3^-/-^ FPUs. These qualitative differences were captured through a 3-point blinded review of MR images (e) and histology images (f). Placental and RPCS volumes were then segmented from all FPUs for both WT and Gab3^-/-^ mice to compare a contrast-to-noise ratio between the higher-signal placental regions and lower signal RPCS and (g) RPCS volumes between WT and Gab3^-/-^ FPUs. Error bars denote standard deviation, significance determined by Wilcoxon rank sum test (* - p<0.05, ** p<0.005).

## Discussion

Gab3^-/-^ pregnant mice experience elevated rates of dystocia that causes stillbirth and lowers the frequency of viable pups as previously reported [18] and is consistent with our observations. Our study was designed to test if high resolution CE-MRI using novel nanoparticle contrast agents, which enable *in vivo* visualization of the RPCS, could be used to detect and characterize the RPCS disruption in Gab3^-/-^ pregnant mouse model of PAS.

Non-contrast T2-weighted anatomical scans in Gab3^-/-^ and WT dams did not visualize the RPCS or any signs of invasion into the myometrial wall. Nanoparticle CE-MRI, however, demonstrated striking differences in FPUs of Gab3^-/-^ pregnant mice. Specifically, the RPCS was thinner and had a more jagged and heterogeneous appearance. Additionally, in many regions, the hypointense RPCS region was either not visible or frankly interrupted. Higher resolution CE-CT confirmed these qualitative findings and allowed better visualization of placental margins and subtle imaging details including scalloping, irregularity in placental shape, and abnormal vascularity. Quantitative assessment of RPCS shape and appearance was performed through volumetric segmentation and blinded review of both high-resolution nanoparticle CE-MRI T1w images and histopathology, confirming the qualitative observations that Gab3^-/-^ RPCS was thinner and less visible compared to WT RPCS.

As demonstrated previously in the Gab3^-/-^ model [18] and in a mouse model of surgical uterine scarring [17], our histological analysis demonstrated morphological differences in the appearance of the Gab3^-/-^ RPCS. Abnormal vascularity and irregular placental shape were also seen in H&E stained images, thus confirming findings from contrast-enhanced imaging. Further, as shown in previous work [17,18], delineation of the retroplacental clear space or myometrial interface was not as obvious in Gab3^-/-^ placentae relative to WT counterparts. In previous work, we defined the normal appearance of this interface as a lower stained region compared to the myometrial and labyrinth regions [20]. In this study, the Gab3^-/-^ RPCS had a denser stain and appeared more heavily vascularized, hence the “heterogeneous” characterization. Blinded review scores for histological data showed a greater difference between Gab3^-/-^ FPUs and WT FPUs than blinded assessment of CE-MRI data. The higher density of staining in histological images was a more obvious finding than disruption in the MR images, possible due to the 2D acquisition and difficulties with orienting the space to better observe differences.

The approach in this study is especially valuable as *in vivo* imaging enabled 3D characterization of RPCS disruption and volume loss. The elevated rates of dystocia and complicated pregnancy in the Gab3^-/-^ model warrant further study in the context of modeling placenta accreta. This study represents an advance in assessing late gestational changes of the RPCS in pregnant Gab3^-/-^ mice and could be used to better model PAS disorders and eventually develop new methods of early diagnosis and intervention.

Several limitations to this study are acknowledged. We observed elevated rates of premature delivery in female Gab3^-/-^ mice and chose E16.5 as our experimental timepoint to ensure consistent analysis between experimental and control groups. Testing at a later timepoint, however, may reveal deeper levels of infiltration and will be considered in future studies. Additionally, testing at earlier timepoints may help determine when RPCS disruption occurs during gestational development. In our breeding study, newborn status was not more thoroughly analyzed beyond simply determining viability. Assessing newborn weight or conditions and more thoroughly characterizing delivery in Gab3^-/-^ female mice may yield greater insight into dystocia or other invasion-related disorders. Future work in this model will also consider stronger methods to translate these findings into clinical practice, particularly in terms of enhancing T2-weighted imaging and looking for common clinical indicators of invasion.

In summary, contrast-enhanced MRI using a liposomal-Gd blood-pool nanoparticle contrast agent enabled clear visualization of placental invasion across the RPCS in a mouse model of PAS. An MRI-based method for visualizing RPCS disruption in a mouse model of PAS will enable the translation of sensitive early detection methods for clinical diagnosis of invasive placentation.

## Acknowledgements

The authors acknowledge the Texas Children’s Hospital Small Animal Imaging Facility (SAIF) for micro-CT imaging and Texas Heart Institute Pathology Core for histology of feto-placental tissues. Also acknowledged is the Cincinnati Children’s Hospital Medical Center and Dr. Helen Jones for providing the Gab3^-/-^ mouse model. Financial support for this study was provided by the National Institutes of Health (NIH) Grant No. R01HD094347.

## Author contributions

Guarantor of integrity of entire study: K.G.B., study concepts/study design or data acquisition or data analysis/interpretation: all authors; manuscript drafting or manuscript revision for important intellectual content: all authors; approval of final version of submitted manuscript: all authors; agree to ensure any questions related to the work are appropriately resolved: all authors; literature research: A.A.B, K.B.G., A.V.A, K.A.F.; experimental studies: A.A.B., K.B.G., L.D., M.S., P.B; statistical analysis: A.A.B., K.B.G.; manuscript editing, all authors.

## References

[1] M.A. Belfort, Placenta accreta, Am. J. Obstet. Gynecol. 203 (2010) 430–439. https://doi.org/10.1016/j.ajog.2010.09.013.

[2] L. Azour, C. Besa, S. Lewis, A. Kamath, E.R. Oliver, B. Taouli, The gravid uterus: MR imaging and reporting of abnormal placentation, Abdom. Radiol. 41 (2016) 2411–2423. https://doi.org/10.1007/s00261-016-0752-5.

[3] B.D. Einerson, J. Comstock, R.M. Silver, D.W. Branch, P.J. Woodward, A. Kennedy, Placenta Accreta Spectrum Disorder: Uterine Dehiscence, Not Placental Invasion, Obstet. Gynecol. 135 (2020) 1104–1111. https://doi.org/10.1097/AOG.0000000000003793.

[4] L.A. Brown, M. Menendez-Bobseine, Placenta Accreta Spectrum, J. Midwifery Women’s Heal. 66 (2021) 265–269. https://doi.org/10.1111/jmwh.13182.

[5] E. Jauniaux, J.C. Kingdom, R.M. Silver, A comparison of recent guidelines in the diagnosis and management of placenta accreta spectrum disorders, Best Pract. Res. Clin. Obstet. Gynaecol. 72 (2021) 102–116. https://doi.org/10.1016/j.bpobgyn.2020.06.007.

[6] Y. Hovav, M. Almagor, Risk of choriocarcinoma from postpartum placental remnants staying for extended times in the uterus, Acta Obstet. Gynecol. Scand. 93 (2014) 720. https://doi.org/10.1111/aogs.12367.

[7] K.E. Fitzpatrick, S. Sellers, P. Spark, J.J. Kurinczuk, P. Brocklehurst, M. Knight, The management and outcomes of placenta accreta, increta, and percreta in the UK: A population-based descriptive study, BJOG An Int. J. Obstet. Gynaecol. 121 (2014) 62–71. https://doi.org/10.1111/1471-0528.12405.

[8] I. Kumar, A. Verma, R. Ojha, R.C. Shukla, M. Jain, A. Srivastava, Invasive placental disorders: A prospective US and MRI comparative analysis, Acta Radiol. 58 (2017) 121–128. https://doi.org/10.1177/0284185116638567.

[9] F.N.Y. Yu, K.Y. Leung, Antenatal diagnosis of placenta accreta spectrum (PAS) disorders, Best Pract. Res. Clin. Obstet. Gynaecol. 72 (2021) 13–24. https://doi.org/10.1016/j.bpobgyn.2020.06.010.

[10] M.A. Maher, A. Abdelaziz, M.F. Bazeed, Diagnostic accuracy of ultrasound and MRI in the prenatal diagnosis of placenta accreta, Acta Obstet. Gynecol. Scand. 92 (2013) 1017–1022. https://doi.org/10.1111/aogs.12187.

[11] J.M. Palacios-Jaraquemada, A. Fiorillo, J. Hamer, M. Martínez, C. Bruno, Placenta accreta spectrum: a hysterectomy can be prevented in almost 80% of cases using a resective-reconstructive technique, J. Matern. Neonatal Med. (2020) 1–8. https://doi.org/10.1080/14767058.2020.1716715.

[12] S. Thiravit, K. Ma, I. Goldman, P. Chanprapaph, P. Jha, D.S. Hippe, M. Dighe, Role of ultrasound and mri in diagnosis of severe placenta accreta spectrum disorder: An intraindividual assessment with emphasis on placental bulge, Am. J. Roentgenol. 217 (2021) 1377–1388. https://doi.org/10.2214/AJR.21.25581.

[13] D.M. Twickler, C.S. Yule, C.Y. Spong, Predicting Placenta Accreta Spectrum: Validation of the Placenta Accreta Index, J. Ultrasound Med. 40 (2021) 2789. https://doi.org/10.1002/jum.15663.

[14] E. Jauniaux, A. Bhide, Prenatal ultrasound diagnosis and outcome of placenta previa accreta after cesarean delivery: a systematic review and meta-analysis, Am. J. Obstet. Gynecol. 217 (2017) 27–36. https://doi.org/10.1016/j.ajog.2017.02.050.

[15] J.L. Bailit, W.A. Grobman, M.M. Rice, U.M. Reddy, R.J. Wapner, M.W. Varner, K.J. Leveno, J.D. Iams, A.T.N. Tita, G. Saade, D.J. Rouse, S.C. Blackwell, Morbidly adherent placenta treatments and outcomes, Obstet. Gynecol. 125 (2015) 683–689. https://doi.org/10.1097/AOG.0000000000000680.

[16] T. Svanvik, A.K. Jacobsson, Y. Carlsson, Prenatal detection of placenta previa and placenta accreta spectrum: Evaluation of the routine mid-pregnancy obstetric ultrasound screening between 2013 and 2017, Int. J. Gynecol. Obstet. (2021). https://doi.org/10.1002/ijgo.13876.

[17] S.D. Burke, Z.K. Zsengellér, S.A. Karumanchi, S.A. Shainker, A mouse model of placenta accreta spectrum, Placenta. 99 (2020) 8–15. https://doi.org/10.1016/j.placenta.2020.06.006.

[18] A. Sliz, K.C.S. Locker, K. Lampe, A. Godarova, D.R. Plas, E.M. Janssen, H. Jones, A.B. Herr, K. Hoebe, Gab3 is required for IL-2-And IL-15-induced NK cell expansion and limits trophoblast invasion during pregnancy, Sci. Immunol. 4 (2019). https://doi.org/10.1126/sciimmunol.aav3866.

[19] A.A. Badachhape, L. Devkota, I. V. Stupin, P. Sarkar, M. Srivastava, E.A. Tanifum, K.A. Fox, C. Yallampalli, A. V. Annapragada, K.B. Ghaghada, Nanoparticle Contrast-enhanced T1-Mapping Enables Estimation of Placental Fractional Blood Volume in a Pregnant Mouse Model, Sci. Rep. 9 (2019). https://doi.org/10.1038/s41598-019-55019-8.

[20] A.A. Badachhape, A. Kumar, K.B. Ghaghada, I. V. Stupin, M. Srivastava, L. Devkota, Z. Starosolski, E.A. Tanifum, V. George, K.A. Fox, C. Yallampalli, A. V. Annapragada, Pre-clinical magnetic resonance imaging of retroplacental clear space throughout gestation, Placenta. 77 (2019) 1–7. https://doi.org/10.1016/j.placenta.2019.01.017.

[21] A.N. Shetty, R. Pautler, K. Ghagahda, D. Rendon, H. Gao, Z. Starosolski, R. Bhavane, C. Patel, A. Annapragada, C. Yallampalli, W. Lee, A liposomal Gd contrast agent does not cross the mouse placental barrier, Sci. Rep. 6 (2016) 27863. https://doi.org/10.1038/srep27863.

[22] A.N. Shetty, R. Pautler, K. Ghagahda, D. Rendon, H. Gao, Z. Starosolski, R. Bhavane, C. Patel, A. Annapragada, C. Yallampalli, W. Lee, A liposomal Gd contrast agent does not cross the mouse placental barrier, Sci. Rep. 6 (2016). https://doi.org/10.1038/srep27863.

[23] K. Ghaghada, Z. Starosolski, S. Bhayana, I. Stupin, C. Patel, R. Bhavane, H. Gao, C. Yallampalli, V. George, A. Annapragada, Contrast-enhanced MRI evaluation of placental margins using a nanoparticle blood pool contrast agent: Pre-clinical testing in a pregnant rat model, Pediatr Radiol. 57 (2017) 60–70. https://doi.org/10.1007/s00247-017-3809-x.

[24] K.B. Ghaghanda, M. Ravoori, D. Sabapathy, J. Bankson, V. Kundra, A. Annapraganda, New dual mode gadolinium nanoparticle contrast agent for magnetic resonance imaging, PLoS One. 4 (2009). https://doi.org/10.1371/journal.pone.0007628.

[25] S. Mukundan, K.B. Ghaghada, C.T. Badea, C.Y. Kao, L.W. Hedlund, J.M. Provenzale, G.A. Johnson, E. Chen, R. V. Bellamkonda, A. Annapragada, A liposomal nanoscale contrast agent for preclinical CT in mice, Am. J. Roentgenol. 186 (2006) 300–307. https://doi.org/10.2214/AJR.05.0523.

[26] S.J. Burke, A. Annapragada, E.A. Hoffman, E. Chen, K.B. Ghaghada, J. Sieren, E.J.R. van Beek, Imaging of Pulmonary Embolism and t-PA Therapy Effects Using MDCT and Liposomal Iohexol Blood Pool Agent. Preliminary Results in a Rabbit Model, Acad. Radiol. 14 (2007). https://doi.org/10.1016/j.acra.2006.12.014.

[27] Z. Starosolski, C.A. Villamizar, D. Rendon, M.J. Paldino, D.M. Milewicz, K.B. Ghaghada, A. V. Annapragada, Ultra high-resolution in vivo computed tomography imaging of mouse cerebrovasculature using a long circulating blood pool contrast agent, Sci. Rep. 5 (2015). https://doi.org/10.1038/srep10178.

[28] H. Ishibashi, M. Miyamoto, H. Shinmoto, S. Soga, H. Matsuura, S. Kakimoto, H. Iwahashi, T. Sakamoto, T. Hada, R. Suzuki, M. Takano, The use of magnetic resonance imaging to predict placenta previa with placenta accreta spectrum, Acta Obstet. Gynecol. Scand. 99 (2020) 1657–1665. https://doi.org/10.1111/aogs.13937.

[29] B. Varghese, N. Singh, R.A.N. George, S. Gilvaz, Magnetic resonance imaging of placenta accreta, Indian J. Radiol. Imaging. 23 (2013) 379–385. https://doi.org/10.4103/0971-3026.125592.

